# 3D synthetic microscaffolds promote homogenous expression of NANOG in mouse embryonic stem cells

**DOI:** 10.1101/2020.09.20.302885

**Authors:** Natalia A. Bakhtina, Madlen Müller, Harry Wischnewski, Rajika Arora, Constance Ciaudo

## Abstract

The development of *in vitro* models, which accurately recapitulate early embryonic development, is one of the fundamental challenges in stem cell research. Most of the currently employed approaches involve the culture of embryonic stem cells (ESCs) on two-dimensional (2D) surfaces. However, the monolayer nature of these cultures does not permit cells to grow and proliferate in realistic three-dimensional (3D) microenvironments, as in an early embryo. In this paper, novel 3D synthetic scaffold arrays, fabricated by two-photon polymerization photolithography, are utilized to mimic tissue-specific architecture, enabling cell-to-matrix interaction and cell-to-cell communication *in vitro*. Mouse ESCs (mESCs) are able to grow and proliferate on these structures and maintain their pluripotent state. Furthermore, the 3D microscaffold arrays are integrated into a microscopy slide allowing the evaluation of the expression of key pluripotency factors at the single-cell level. Comparing 2D and 3D surfaces, mESCs grown in serum+LIF on 3D microscaffolds exhibit a stronger and more homogenous expression of NANOG and OCT4 pluripotency factors, than cells cultivated in 2i media, demonstrating that 3D microscaffolds capture naive pluripotency *in vitro*. Thus, the slide affords a novel and unique tool to model and study mammalian early development with greater physiological relevance than conventional 2D cultures.

## 1. Introduction

*In vivo*, cells are arranged in a three-dimensional (3D) multifunctional environment. The chemical and mechanical properties of this specific entourage influence their intracellular functioning. However, most of the currently employed approaches in cell-and tissue-based engineering studies still involve two-dimensional (2D) surfaces, or monolayer cell cultures, that offer unnatural growth kinetics and cell attachments.^[1,2]^ On 2D cultures, cells adhere to a plate, which restricts them to a flat shape. Moreover, these cultures lack a complex tissue-specific architecture leading to dramatic variation in the diffusion of chemical cues and cell-to-cell/cell-to-matrix interactions.^[2]^ As a result, the molecular pathways that regulate cell behaviors are also altered, leading to distinct cellular phenotypes. Recent advances in cell biology, microfabrication techniques and tissue engineering have enabled the development of a wide range of 3D cell cultures, including multicellular spheroids, organoids, scaffolds, hydrogels and 3D bioprinting.^[3]^ Nowadays, they have become a promising alternative for bridging the gap between *in vitro* cultures and living tissues as they exhibit protein expression patterns and intracellular junctions that are more similar to *in vivo* states.^[4]^

Currently, hydrogels are the most widely used system for 3D cell cultures to mimic extracellular matrix *in vitro*. For example, Matrigel, which is reconstituted from the mouse sarcoma and composed of laminin, entactin, collagens and heparin sulfate proteoglycan plus an unknown mixture of growth factors and enzymes, is commonly used in biology.^[5,6]^ It is important to understand that some properties of the hydrogels have major drawbacks, such as very low rigidity, which does not mimic the naturally stiff environment of some tissues.[7] To ensure optimal performance, swelling of these 3D biomaterials, their permeability to different molecules, interaction with media and the immobilization of biomolecules have to be optimized.^[5]^ Since culture composition is not clearly defined, these matrices lead to relatively poor reproducibility and lack of consistency between batches. Finally, the bigger or more complex the 3D volume is, the more difficult the extraction of cells for further experimentations becomes. All these limitations make hydrogels unsuitable for tissue modeling *in vitro*.^[6]^ Consequently, toward the development of scalable 3D culture system, cost-effective, chemically defined, and reproducible culture systems are required.

3D synthetic microscaffolds, fabricated by two-photon polymerization (2-PP) photolithography, offer favorable cell responses due to tunable chemical, physical and mechanical properties (reviewed by Hippler *et al*.).^[8]^ 2-PP technology allows the fabrication of volumetric structures of arbitrary shape by directly writing the intended geometry within a photosensitive material. Due to the unprecedented flexibility of 2-PP, matrix architecture and pore size can be controlled with a resolution down to 100 nanometers.^[9]^ Indeed, structures with well-engineered nanotopographies have features on similar length scales to cellular components and have become useful tools for controlling the cellular environment.^[10]^ For example, artificial surface nanotopography has been shown to maintain long-term human ESC pluripotency by inhibiting cell spreading, which makes the cells less flat, thus increasing the clone integrity.^[11]^ In addition, 2-PP-fabricated scaffolds deliver cells with spatially defined adhesion sites and can instruct different cell types with respect to their proliferation and migration (e.g. by biological functionalization with several proteins).^[12]^ Chemically defined media combined with the 3D architectures, which more accurately resemble the extracellular environment, offer a powerful tool to mimic specific tissues *in vitro*. The employment of necessary mechanical cues would have the important advantage of minimizing the use of biochemical molecules, which are otherwise necessary to regulate cellular phenotype.^[13,14]^

MESCs are a broadly used experimental model system to understand early mammalian development.^[15,16]^ They are derived from the inner cell mass (ICM) of pre-implantation embryos and can be maintained in a pluripotent state *in vitro* using specific cues.^[17]^ Two main pluripotent states have been described *in vitro*: the “naïve” state, which corresponds to blastocyst at days 3.5 (E3.5) – day 4.5 (E4.5) *in vivo* – and the “primed” state, which occur at a later developmental stage E5.5 – E6.5.^[18,19]^ These two states are maintained in two different culture media *in vitro*: naïve in “2i” medium (for two small molecule inhibitors PD0325901 (PD) and Chir99021 (CH)) supplemented with LIF (leukemia inhibitory factor) and primed in serum supplemented with LIF (see details in the Experimental Section).^[18]^ Although both naïve and primed mESCs have the potential to form all somatic cell lineages as well as germ cells, they are distinct in their morphological, epigenetic and transcriptional characteristics.

For example, naïve mESCs grown in 2i form compact dome shaped colonies, whereas primed mESC colonies grown in serum+LIF are larger and flatter in shape.^[20]^ The gene expression program of stem cells is maintained through the action of three key pluripotency transcription factors: OCT4, SOX2 and NANOG. In serum+LIF, mESCs express the NANOG protein in a heterogeneous manner,^[21,22]^ whereas in 2i medium its expression is more homogenous.^[23]^ Indeed, the heterogeneity of NANOG in the stem cells population can be used to distinguish naïve and primed mESCs.

In recent years, a number of experimental strategies and computational models have been applied to reveal the molecular mechanisms and interactions that orchestrate pluripotency (reviewed by Martello *et al*. and Young *et al*). ^[19,24]^ These strategies and models are based mainly on 2D cell cultures, in which cellular phenotypes are regulated by selective suppression and/or activation of key signaling pathways using growth factors and small molecules. Defining an optimal 3D model that best mimics the specificity of the *in vivo* microenvironment is a crucial step towards generating data that accurately reflects what occurs in embryos.^[25]^ Several reports suggest that the critical signals for pluripotency maintenance likely depend more on spatial conformation changes rather than on extrinsic growth factors.^[14,26]^ For example, Nava *et al*. used highly resolved “nichoid” in order to compare the expression of pluripotency and differentiation markers induced in mESCs, thereby showing the potential of 3D cultures for reducing the use of biochemical molecules.^[13]^ However, none of the previous studies were able to distinguish between different mESC populations – naïve and primed – based on their pluripotent potential.

In this paper, we presented several designs of tailored 3D microscaffold arrays using 2-PP photolithography and evaluated their impact on mESC pluripotency. We demonstrated that these microscaffolds maintained mESCs in a pluripotent state. In comparison to a 2D solid film, mESCs in 3D microscaffold arrays exhibited a stronger signal intensity of two pluripotency markers – NANOG and OCT4. Tracking the heterogeneity of the NANOG pluripotency factor by fluorescence microscopy allowed us to demonstrate that 3D solid culture reinforce naïve pluripotency of mESCs.

## 2. Results and Discussion

### 2.1 3D synthetic microscaffold array designs

The key aspect in the design of 3D microscaffold arrays was to mimic the complex multifunctional environment required for promoting mESC adhesion and proliferation. Therefore, the design of the microscaffolds was motivated by the following reasons. It should: (1) be of highly precise micrometer shape (i.e. scale comparable to the cell size) with a roughness on the nanometer level (down to 100 nm) to allow for highly repetitive motifs for cells; (2) have a structural rigidity (as mechanical support); (3) be able to physically contain mESCs within their 3D microarchitecture, like individual niches (since mESCs have a round shape, microscaffolds with round/square form were prefered over rectangular/triangular microscaffolds); (4) have an open access geometry of 3D microscaffolds, which would allow the optical access; (5) allow for spatial cell distribution by gravity driven sedimentation into individual microscaffolds; (6) guarantee cell-to-matrix interaction in a 3D environment; (7) provide homogeneous dispersion of nutrients and chemical cues as well as gas exchange.

In order to evaluate the impact of scaffold architectures on mESC culture, we designed several concepts, including ring-like (15 µm × 15 µm × 10 µm in terms of width × length × height), basket-like (15 µm × 15 µm × 10 µm in terms of width × length × height) and post-like (2 µm in diameter and 2 µm distance between posts). **Figure 1A** schematically shows the three microscaffold designs. Ring-like and basket-like microscaffold designs allow for the allocation of each cell to an individual microscaffold. In post-like microscaffolds, cells can hardly migrate due to a dimensional incompatibility enabling the planar cell distribution.

**Figure 1.**
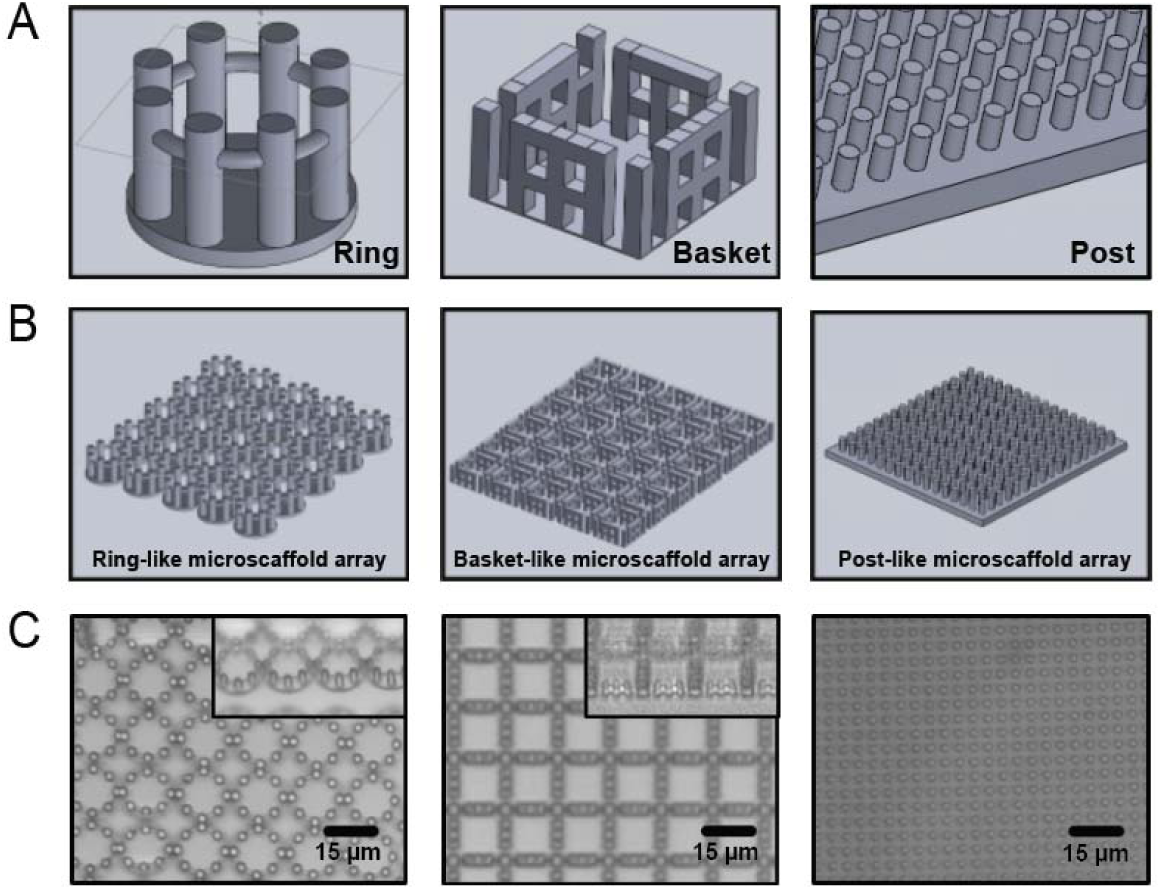
3D microscaffold arrays. (A) Schematic views of three 3D microscaffold designs, including ring-like, basket-like and post-like. (B) The corresponding schematics of 3D microscaffold arrays. (C) Photographs of the 3D microscaffold arrays (top view) fabricated by two-photon polymerization (2-PP) photolithography on a glass coverslip (digital microscope, Keyence, Japan). The total surface area of the array is 2.25 mm^2^. The insets are a photograph of 3D microscaffolds taken at an angle of 45°.

To recapitulate the cell-to-cell communication network, these structures were then assembled into tightly packed arrays. The number of microscaffolds in the array was motivated by a relatively large surface array (i.e. millimeter range) in order to perform cell population analysis; and should guarantee cell-to-cell interaction (i.e. cells in the neighboring scaffolds). Therefore, individual scaffolds have to be in direct contact. Besides the architectural aspects, the design of both the 3D microscaffolds and the array must also satisfy technological limitations of the 2-PP process. For example, high aspect ratio structures (1:10) should be avoided because microstructures might collapse after the development step. Large surface area elements (i.e. centimeter range) are also undesirable at the cost of a corresponding increase in overall fabrication time. The schematics and photographs of 3D microscaffold arrays are depicted in **Figures 1B** and **1C**, correspondingly.

### 2.2 Culture of mESCs in 3D microscaffold arrays

To evaluate the biocompatibility of 3D microscaffolds (i.e. cell viability, adhesion to the 2PP-engineered microscaffolds and morphological compatibility), we first fabricated all three 3D microscaffold arrays on a glass coverslip (see “Experimental section”). After coating of 3D microscaffold arrays with gelatin, mESCs were seeded and cultured under defined serum+LIF conditions for 5 days. The morphology of mESCs was evaluated daily by live microscopy. As an example, we show the mESCs grown in the ring-like microscaffold array in **Figure 2**. Rapidly growing colonies were observed from days 1 to 5 suggesting that the 3D microscaffold arrays can maintain the growth of mESCs under serum+LIF condition.

**Figure 2.**
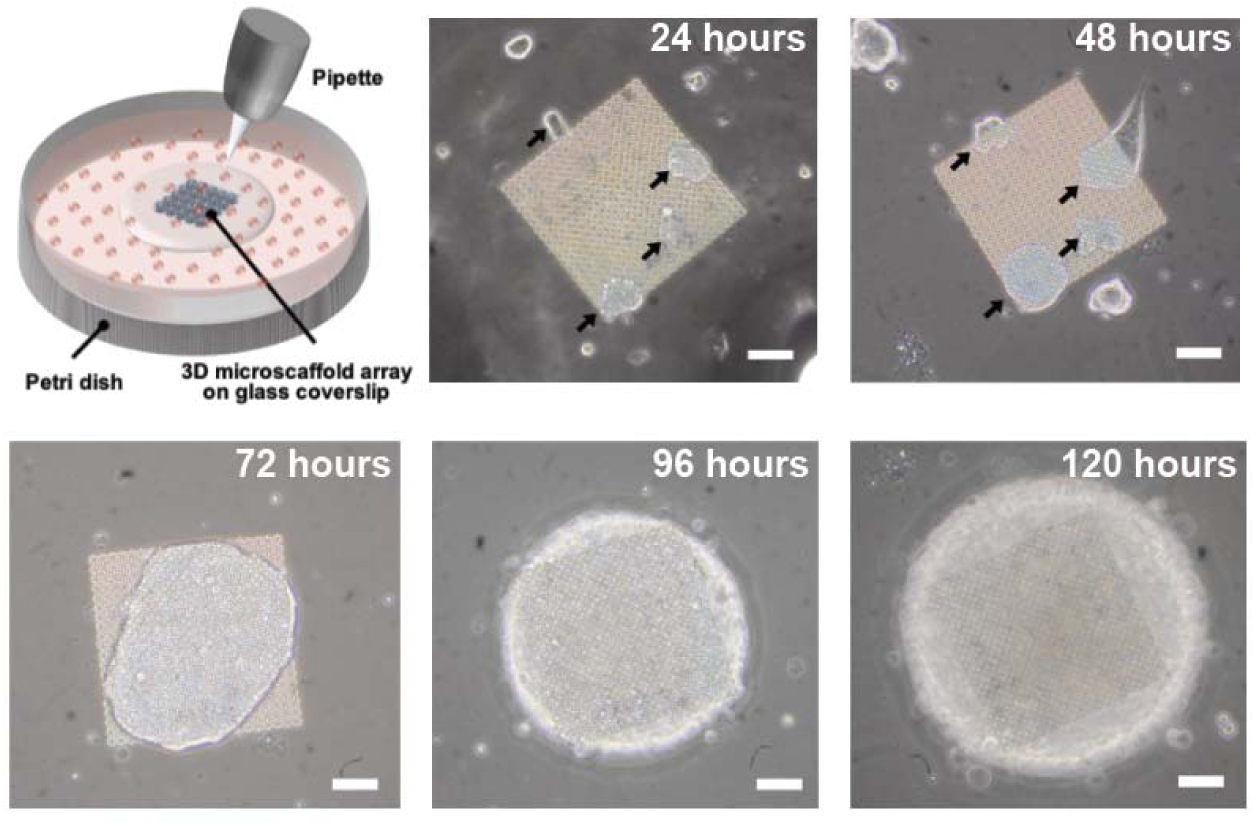
Schematic view of mESC seeding procedure on the ring-like microscaffold array (total surface area of the array is 400 μm^2^) and time-lapse phase contrast images (24 hours – 120 hours). After day 1 cells adhered extensively to the microscaffolds, after 2 days – formed individual cell aggregates on the top of the 3D microscaffold array and after 3 days – proliferated to fused round-shaped colonies typical for mESCs. Cell colonies are highlighted by black arrows. The scale bar is 50 µm.

We then assessed the stemness properties of the mESCs grown on the 3D microscaffold arrays using alkaline phosphatase (AP) staining.^[20,23]^ Cell populations were maintained in all three 3D microscaffold arrays (ring-like, basket-like and post-like) under serum+LIF conditions for 3 days. They formed distinct AP-positive colonies and the quantification of integrated AP intensity at the single colony level are shown in **Figure 3**. To initiate differentiation on 2D surface (used as a control), mESCs were cultured on glass coverslips in LIF-free media (see “Experimental section”). In this differentiation media, cells did not form colonies resembling embryonic stem cells and had less AP intensity (highlighted by light magenta color), indicating that these cells committed to differentiation. The quantification of the single colonies revealed higher AP intensity for all three 3D microscaffold arrays compared to mESCs on 2D surface. These initial experiments convincingly demonstrated that mESCs retained their self-renewal potential and remained undifferentiated in the 3D microscaffold array under serum+LIF conditions.

**Figure 3.**
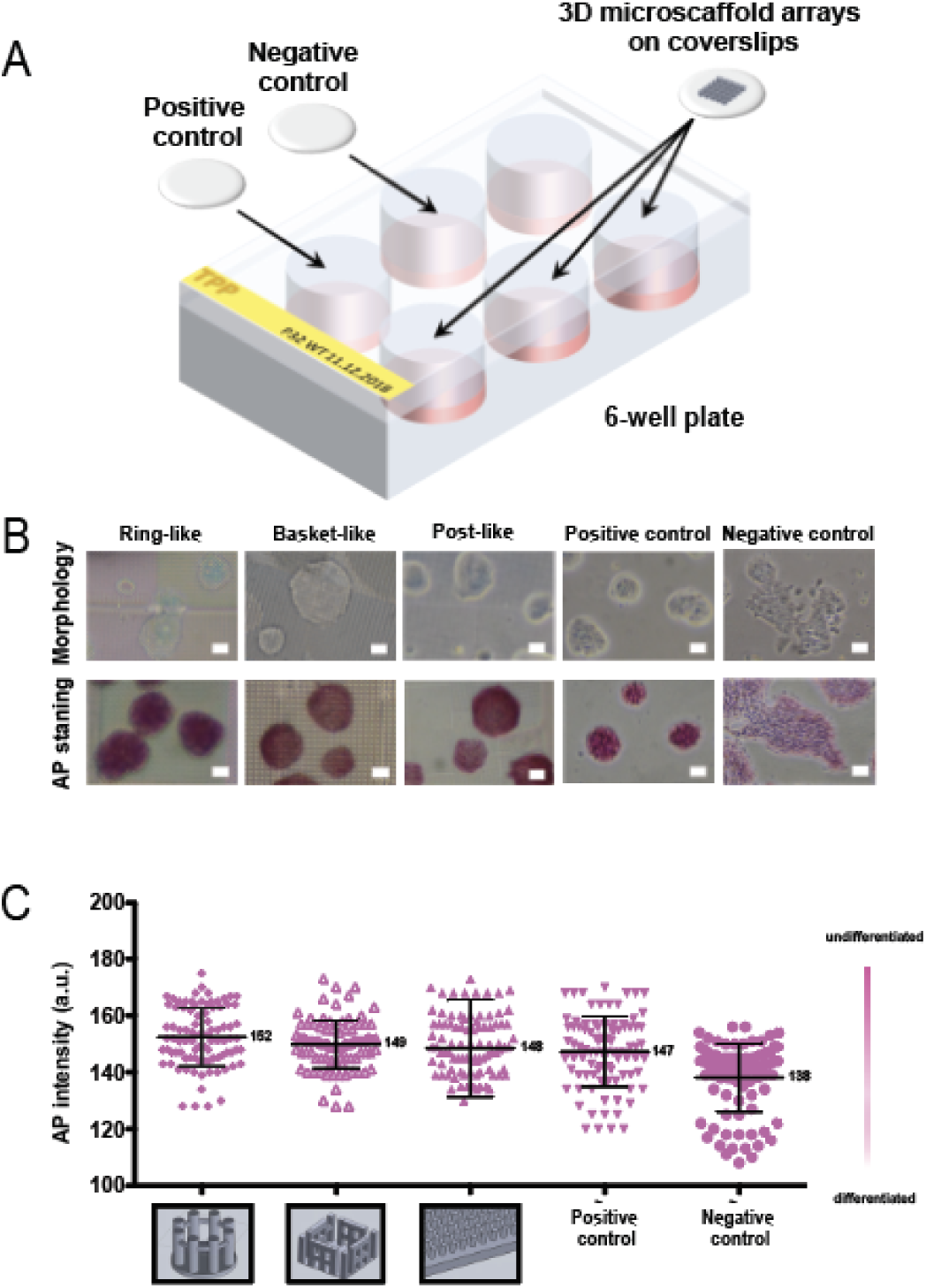
Schematic view of the experimental design used for alkaline phosphatase (AP) staining (A). AP-stained mESC colonies cultured in three 3D microscaffold arrays (ring-like, basket-like and post-like) and on the glass coverslips (B). Positive control: mESCs were maintained in serum+LIF medium. Negative control: mESCs were cultured in differentiation medium (see “Experimental section”). Cell colonies were imaged 72 hours after seeding. The scale bar is 50 µm. Quantification of AP intensity in mESCs at the single colony level (C). Quantification of AP stained colonies from three independent experiments.

### 2.3 3D synthetic microscaffolds promote NANOG homogenous expression

In order to assess the pluripotent state of the mESCs grown in 3D microscaffolds at the single cell level, we integrated 3D microscaffolds on a diagnostic microscopy slide (**Figure 4A**). Quantitative immunofluorescence analysis provides information, which includes location and distribution of protein levels in single cells across populations or relative amounts of two or more proteins within a single cell.^[22,27,28]^ The use of the slide significantly simplified the immunofluorescence imaging, since it was supplied with a thin pre-patterned hydrophobic epoxy resin mask. Firstly, the mask enabled handling of several cell populations in parallel and subject each population to an experimental protocol in a systematic manner while decreasing reagent consumption (see “Experimental section”). Secondly, it protected the microscaffold arrays from damage during preparation of the slide for immunofluorescent imaging by eliminating the direct contact of the coverslip and microscaffolds. And finally, it enabled high resolution imaging by providing the short working distance from the microscope objective to 3D microscaffolds.

**Figure 4.**
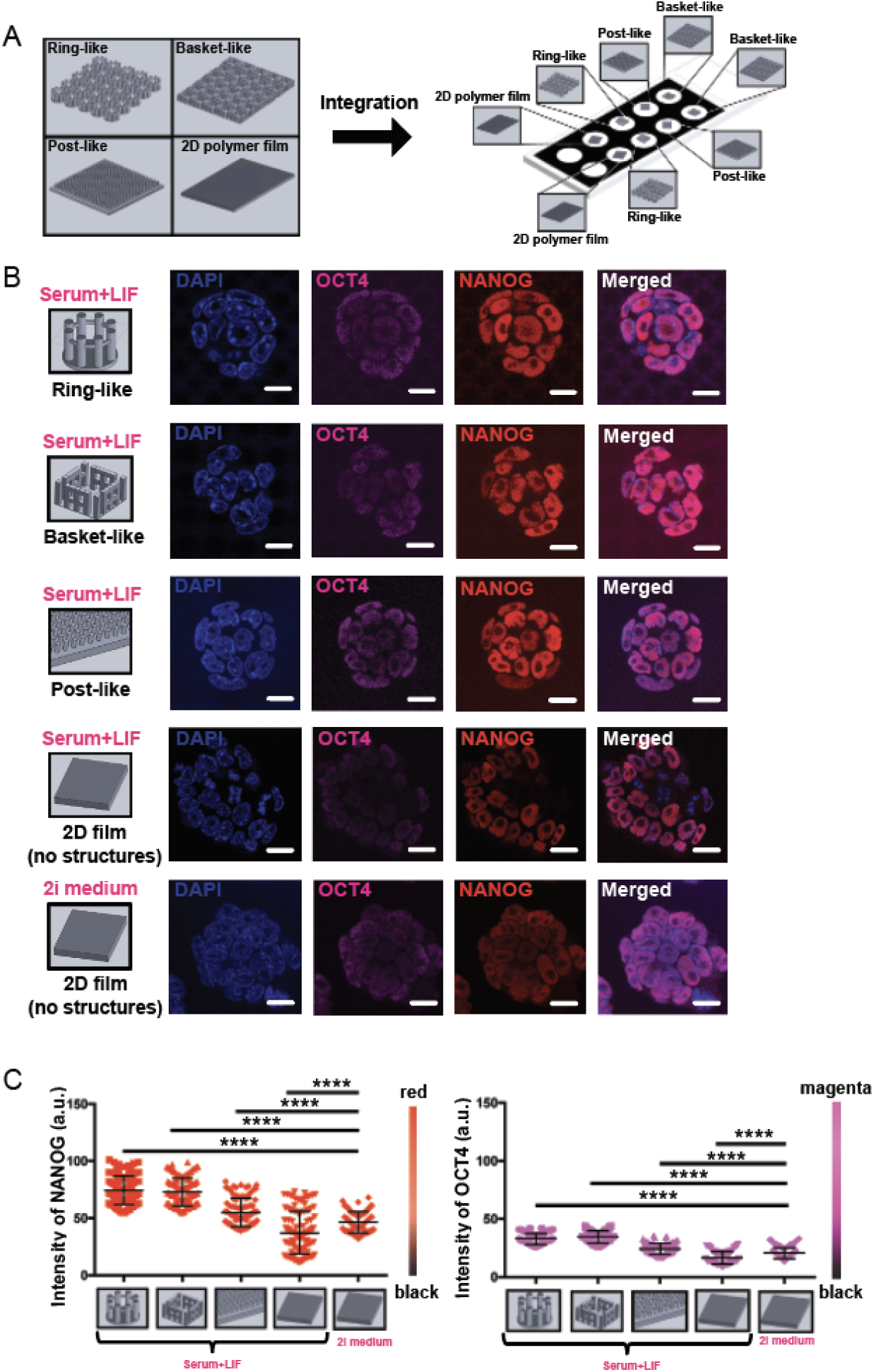
Schematic views of three 3D microscaffold arrays (ring-like, basket-like and post-like), which were fabricated by two-photon polymerization (2-PP) photolithography in the wells of the diagnostic microscopy slide (A). The 2D solid polymer film was used as a control. Representative immunofluorescent images of NANOG (red) and OCT4 (magenta) expression in mESCs grown 3D microscaffold arrays and on 2D film in serum+LIF (B). As a control, cells were maintained on 2D film in 2i medium. DAPI was used as a nuclear counterstain. Scale bar is 10 µm. Quantification of immunofluorescent intensity for the NANOG (red) and OCT4 (magenta) at the single nuclear level in mESCs cultured in three 3D microscaffold arrays (ring-like, basket-like and post-like) and on 2D solid polymeric film under serum+LIF and 2i conditions (C). Number of counted nuclei n=100. Immunofluorescence staining was performed in triplicate. P<0.0001 (Mann–Whitney test).

To investigate the effects of 3D microscaffolds on the shape of stem cell colonies, mESC populations were maintained in all three 3D microscaffold arrays (ring-like, basket-like and post-like) under serum+LIF conditions for 48 hours. In the basket-like microscaffold array, cells dropped into individual microscaffolds, settled down, adhered to the scaffold walls and grew in height. In the ring-like microscaffold array, cells occupied not only the volume within the microscaffolds, but also the space in between (**Figure S1A**, Supporting Information 1). MESCs adopted the form of the given internal volume of both the ring-like and basket-like microscaffold arrays and showed structural reorganization of subcellular microarchitecture. When cells were grown on the post-like microscaffold array and 2D layer (i.e. solid polymer film), they were flatter and did not reveal any defined axis formation or shape. MESCs have also demonstrated a tendency to form diverse colonies within the 2PP-engineered microscaffolds. The Z-stack side-view projections of confocal images acquired on the microscaffold arrays showed that mESCs formed round-shaped colonies. A side-view projection of the ring-like microscaffold array is depicted in **Figure S1B** (Supporting Information 1) as an example. These results imply that the microscaffolds are able to guide the spatial organization of colony formation, thus limiting the need for selective cell seeding in stem cell culture systems. We hypothesize that the ability of mESCs to grow inside the 2-PP engineered microscaffolds better mimic the *in vivo* state, where cells are allowed to self-organize in a truly 3D multifunctional environment.

To probe the dynamics of the pluripotent states and examine whether the 3D architecture of microscaffolds rather than the material itself impacts expression levels of pluripotency factors, we compared mESC populations in all three microscaffold arrays and on 2D solid polymer film under serum+LIF and 2i conditions. An optimized protocol for immunostaining is presented in **Figure S2**, Supporting Information. The expression of OCT4 and NANOG proteins was monitored at the single cell level. In agreement with previous studies, the distributions of expression values (black bars) of the two pluripotency markers – OCT4 and NANOG – on 2D solid film across the population were representative for each growth condition.^[20–22,29]^ Representative immunofluorescent images of individual cellular nuclei are shown in **Figure 4B**. In serum+LIF, NANOG protein had a high degree of heterogeneity, which is shown by a relatively broad distribution of expression values (black bars in **Figure 4C**, serum+LIF conditions). In contrast, OCT4 did not exhibit heterogeneity. As in pre-implantation embryos, OCT4 is present in all cells of the inner cell mass (ICM) until late blastocyst.^[30]^ NANOG heterogeneity among the cell populations was reduced on 3D microscaffolds, which is shown by a relatively narrow distribution of expression values (black bars in **Figure 4C**, serum+LIF conditions). Upon the transfer into 2i conditions, mESCs in 3D microscaffold arrays and on the 2D solid film showed significantly higher mean expression levels of both pluripotency markers (**Figure 4C**). Moreover, the rather homogeneous expression levels of the NANOG transcription factor in 3D microscaffolds indicated that the cell populations possessed a stronger self-renewal ability (figure panel **Figure 4B**, 2i conditions). Interestingly, mESCs cultured in all 3D microscaffold arrays in serum+LIF conditions expressed NANOG and OCT4 at levels comparable to those values in 2i medium (**Figure S3**, Supporting Information 3). These results indicated that mESCs show an enhanced responsiveness both to 3D microscaffolds and 2i medium conditions. It can be hypothesized that mESCs in 3D microscafold arrays might have a stronger self-renewal ability resembling more closely the ICM of pre-implantation embryos.^[23,31]^ Therefore, the physical properties of 3D microscaffolds arise from their patterned structure, rather than an innate property of the material. Changing the constituent material or the surface chemistry may also impact the response (as is true for physical metamaterials), but the main effect is derived from the structure. Besides the 3D environment, the nanotopography of the 3D microscaffold’s surface, as a result of the ellipsoidal two-photon absorption volume, may also amplify the expression levels of pluripotency markers.^[32,33]^ Our data demonstrated that in comparison to 2D surface, mESCs in 3D microscafold arrays exhibit a stronger immunostaining signal of two pluripotency markers – NANOG and OCT4 – and show a more homogenous pluripotency state as highlighted by the expression of NANOG.

## 3. Conclusions

We present a controllable engineered-niche system for studying the biophysical regulation of stem cell pluripotency. Revealing the molecular basis of mESC cellular heterogeneity in 3D culture conditions is not only important for understanding the flexible nature of the pluripotent state but might also serve as a model to understand heterogeneity in other systems (e.g. human ESCs and induced pluripotent stem cells (iPSCs)). The design of tailored 3D microscaffolds and the array dimensions can be easily adapted to any cell type. Due to the unprecedented flexibility of the 3D patterning approach and superior advantages in terms of material characteristics, the entire manufacturing process can be performed in a time-efficient manner. Thus, the technology is widely applicable to study other biological systems, for which 3D environment is of crucial importance for proper functioning.

## 4. Experimental Section

### Design and manufacturing of 3D microscaffold arrays

A 3D computer-aided design program (Solidworks Corp., USA) was used for 3D model development of the microscaffolds. An original file (*.sldprt) of a 3D solid object was converted into Surface Tesselation Language (STL) file for the “Photonic Professional GT“system (Nanoscribe GmbH, Germany). Two software packages, DeScribe 2.2.1 and NanoWrite 1.7.6, were used to control the system. To define the structural design, the input file (STL) was converted by DeScribe 2.2.1 to the Photonic Professional’s native data format (GWL), where the number of microscaffolds in the arrays as well the fabrication parameters, such as laser power, laser power scaling, line distance and scanning speed of the laser focus, were configured. The generated GWL file with a code was then loaded for direct laser writing process by the control software NanoWrite for manufacturing. An example of a GWL code segment can be found in Supporting Information 4.

The manufacturing of 3D microscaffold arrays was performed on a glass coverslip and diagnostic microscopy slide with a pre-patterned hydrophobic epoxy resin mask (the thickness may vary between 30 µm and 50 µm) with 10 reaction wells. Prior to patterning, the glass coverslip and slide were washed with acetone and isopropanol to clean the surface and increase the hydrophilicity of the surface before a drop of the photosensitive material (IPL-780 photoresist, Nanoscribe GmbH, Germany) was placed on the top. One droplet (a volume of a few microliters) was deposited manually on the glass coverslip or in the fluidic reservoirs of the diagnostic microscope slide using a pipette. Afterwards, the coverslip/slide was fixed to the sample holder by a gluing tape and placed in a holder that fits into the piezoelectric stage. Microscaffold arrays (one microscaffold array per coverslip or well of the diagnostic slide) were written in a “bottom-up” sequence (i.e. the first layer was attached on the substrate surface). To minimize the optical aberrations related to the immersion-oil configuration and then get the best results in terms of resolution, the photoresist-immersion configuration (Dip-in Laser Lithography, DiLL) was used. The writing speed was 40 000 μm s^−1^ for achieving completely crosslinked polymeric structures with well-defined 3D geometry and structural rigidity. To ensure that the polymerized material had a good connection to the substrate and to enhance the mechanical stability, the writing volume overlapped a few micrometers with the substrate. During the post-treatment, the uncured material was removed in a two-step development process: (1) 5 minutes in mr-DEV600 (Micro Resist Technology GmbH); (2) 15 minutes in 2-propanol. Finally, the substrate was dried with nitrogen gas. The 2D solid film (i.e. 170 µm thick macro-controlled pattern), which is used as a 2D control, was fabricated by placing a droplet (∼ 10 µL) of the same photoresist on a microscope slide between two quartz glass slides (170 µm thick). A quartz glass slides was placed on top to ensure a homogeneous photoresist distribution with no trapped air bubbles in the liquid photoresist. The microscope slide with liquid photoresist was then cured under UV-light (365 – 405 nm) for 2 minutes (MA6, Karl Süss, Germany).

### Cell culture

The mESC line used in this study was E14Tg2A (CRL-1821, ATCC). In serum+LIF conditions, cells were cultured in DMEM (Invitrogen) supplemented with 15 % fetal bovine serum (Gibco), 1 μL/mL LIF (EMD Millipore), 0.1 mM 2-β-mercapto-ethanol (Thermo Fisher Scientific), 0.05 mg/mL streptomycin, and 50 μL/mL penicillin (Sigma-Aldrich). In differentiation medium (i.e. LIF-free conditions), cells were cultured in DMEM (Invitrogen) supplemented with 10 % fetal bovine serum (Gibco), 0.1 mM 2-β-mercapto-ethanol (Thermo Fisher Scientific), 0.05 mg/mL streptomycin, and 50 μL/mL penicillin (Sigma-Aldrich). In 2i conditions, cells were cultured in N2B27 buffer (Cellartis) complemented with 50 U/mL of penicillin and 0.05 mg/mL of streptomycin and the following inhibitors: PD0325901 (Millipore) at 1 μM final concentration, CHIR99021 (StemCell Technologies) at 3 μM final concentration and 1000 U/mL of LIF.

### Biocompatibility test of microscaffold’s material

Before cell seeding, the glass coverslip (25 cm in diameter) with integrated 3D miroscaffold array was washed with 70 % ethanol and sterilized by UV-light (2.5 hours). Afterwards, it was coated by 0.2 % gelatin (1 hours) and placed in a Petri dish (4 cm in diameter). The cells were suspended in standard conditions (serum+LIF) and seeded at a density of 16 000 cells/cm^2^. Media was exchanged after 24 hours. All cells were grown at 37 °C in 8 % CO_2_ from day 1 to day 5 and analyzed continuously by bright field microscopy (Nikon Eclipse TS100).

### Alkaline phosphatase (AP) staining

Glass coverslips (13 cm in diameter) with or without integrated 3D microscaffold arrays were washed with 70 % ethanol, placed in the 6-well plate, sterilized by UV-light (2.5 hours) and then coated by 0.2 % gelatin (1 hour). The cells were suspended in standard conditions (serum+LIF medium) and seeded at a density of 9 500 cells/cm^2^. Media was exchanged after 24 hours. The alkaline phosphatase (AP) staining was performed 72 hours after seeding using the Leukocyte Alkaline Phosphatase kit (Sigma, St. Louis, MO, USA) according to the manufacturer’s instructions. AP staining images were captured by microscopy (Nikon Eclipse TS100). Image analysis was performed using Fiji software (https://imagej.net/Fiji). Integrated intensity measurements for the AP-stained mESC colonies were obtained after delineating the colony as the region of interest for segmentation. Only accurately segmented colonies were included in the analyses. 100 colonies from three independent experiments were analyzed for each sample. Scatterplots and statistical analysis of the data were generated using Prism 6 (https://www.graphpad.com/scientific-software/prism/). Mann-Whitney test was used to compute statistical significance. P-value of p < 0.05 were considered to be significant.

### Immunostaining

Before cell seeding, the 10-well diagnostic microscopy slide with integrated 3D miroscaffold arrays were washed with 70 % ethanol and sterilized by UV-light (2.5 hours). Afterwards, it was placed in a standard grade plastic vessel with 3–4 mL of phosphate-buffered saline (PBS×1) buffer to avoid medium evaporation. 3D microscaffold arrays and 2D polymeric film were then coated by 0.2 % gelatin (24 hours). Cell seeding was performed in two steps: (1) placing a 50 μl medium droplet per well; (2) injecting 1 μl with ∼100 cells above polymer foil or 3D microscaffold array. Media was exchanged after 24 hours. All cells were grown at 37 °C in 8 % CO_2_ from day 1 to day 2 and analyzed 48 hours after seeding.

Indirect immunofluorescence (IF) was performed 48 hours post seeding. Cells grown on microscaffolds were washed with 1x PBS and fixed in 3.7 % formaldehyde for 10 minutes. Cells were next washed in 1x PBS three times and permeabilized with CSK buffer (100 mM NaCl, 300 mM sucrose, 3 mM MgCl_2_, 0.5% Triton X-100, 10 mM PIPES, pH 6.8) for 5 minutes on ice. Cells were blocked in 2.5 % BSA, 0.1 % Tween 20 in 1x PBS for 1 hour at room temperature followed by incubation with the primary antibody diluted in the blocking solution for 1 hour at room temperature. The following antibodies were used NANOG antibody (D2A3, Cell Signaling, 1:500) and OCT4 antibody (611202, BD Biosciences, 1:500). Cells were washed three times in 1xPBS with 0.1 % Tween 20 prior to incubation with secondary antibodies, Donkey anti goat conjugated with Alexa Fluor 568, Invitrogen, 1:4000) and Donkey anti mouse conjugated with Alexa Fluor 647 (Invitrogen, 1:4000). Cells were washed three times in 1xPBS with 0.1 % Tween 20 and once in 1xPBS, counterstained with 4’,6-diamidino-2-phenylindole (DAPI) (100µg/mL) for 10 minutes. Vectashield® (Antifade Mounting Medium) was used to reduce photobleaching.

### Imaging and image analysis

Using sequential scanning mode images were acquired with Leica SP8-AOBS-CARS laser confocal microscope equipped with a 40x 1.4 NA water HC PL APO CS2 objective. The same imaging parameters, such as laser intensity, gain, and pinhole, were set for all samples. Range indicator palette option was used to ensure that no oversaturated images were taken. Imaged colonies were randomly selected. Autofluorescence of the material was evaluated with a multiphoton (MP) confocal microscope (Leica SP8 MP, Leica Microsystems, Germany). In particular, we observed more autofluorescence of the photoresist in the blue and green region of the emission spectrum and less autofluorescence in the far-red region. To this end, excitation of the fluorophores conjugated to the secondary antibodies was implemented with wavelengths above 500 nm. Confocal images from an optical section located 1–2 μm from the top of the 3D microscaffold arrays were saved independently in grayscale for each of the channels. Due to the autofluorescence of the 3D microscaffold arrays, cells positioned at the beginning (i.e. at the bottom of 3D microscaffolds) and end of the z stack (i.e. top layer of cell colony) were excluded from the analysis. Image analysis was performed using Fiji software (https://imagej.net/Fiji). Briefly, integrated intensity measurements for the red (NANOG at 567 nm emission wavelength) and far red (OCT4 at 647 nm emission wavelength) channel were obtained after delineating the nucleus as the region of interest using DAPI staining for segmentation. Only accurately segmented nuclei were included in the analyses. 100 nuclei from three independent experiments were analyzed for each sample. Scatterplots and statistical analysis of the data were generated using Prism 6 (https://www.graphpad.com/scientific-software/prism/). Mann-Whitney test was used to compute statistical significance. P-value of p < 0.05 were considered to be significant. The quantitative immunofluorescence analysis was based on a number of assumptions: (1) the antibody is specific to the antigen, thus a blocking step was included in the protocol to avoid unspecific binding of the antibody to the antigens; (2) the antibody binds to all available specific antigens, consequently a permeabilization step was included in the protocol to allow the access of the antibody to the antigens through nuclei membrane and increase its chances of binding; (3) the intensity of the fluorescent signal is proportional to the concentration of the antigen; (4) there might be variability across samples, therefore to reduce this variability, when different cell lines were tested for their OCT4 and NANOG intensities, the whole experiment was performed in parallel including cell seeding, the complete protocol for fluorescent immunohistochemistry and imaging steps.

## Supporting Information

Supporting Information is available from the Wiley Online Library or from the author.

## Acknowledgements

We thank the Ciaudo lab for the critical reading of the manuscript and for fruitful discussions. This work was supported by a core grant from ETH-Z (supported by Roche) and Olga MayenFisch Stiftung. N.A.B. is supported by the European Union (EU) [Marie Sklodowska-Curie Individual Fellowship (project 792776, 8th Research Framework Programme HORIZON 2020). We also express our gratitude to the Air Quality Control Laboratory, ETH Zurich (in particular, to Jean Schmitt) and the micro-and nanofabrication facilities at Binnig and Rohrer Nanothechnology Center (BRNC), ETH Zurich for providing access to the equipment for 3D printing and assistance in 3D microscaffold manufacturing. We are also thankful to the Scientific Centre for Optical and Electron Microscopy (ScopeM, ETH Zurich) for support provided in high resolution intracellular imaging.

## Author Contributions

N.A.B. and C.C. wrote the manuscript. N.A.B designed and fabricated the 2D microscaffold arrays. N.A.B assembled all the figures. Biological experiments were supported by M.M, RA and HW. All authors read and approved the final manuscript.

