## Supplemental Material for "3D synthetic microscaffolds promote homogenous expression of NANOG in mouse embryonic stem cells"

**Supporting Information 1**

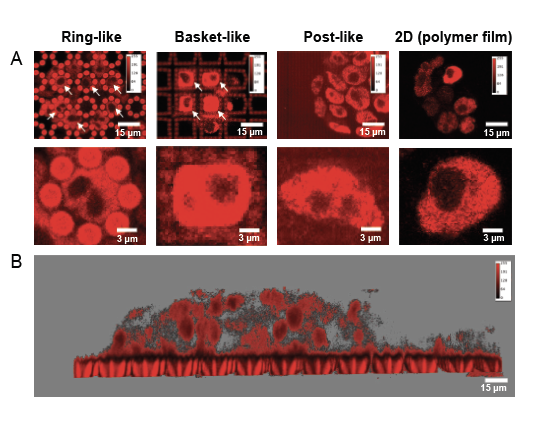

**Figure S1**. Immunofluorescent images showing the expression of NANOG (red) in mESCs. (A) Three microscaffold arrays, including ring-like on the left, basket-like in the middle and post-like on the right (total surface area of the array is 2.25 mm^2^), are visualized on subsequent horizontal section located at the arrays' middle. (B) Z-stack side-view projection of immunofluorescent images acquired on the ring-like microscaffold array. The 2D solid polymer film was used as a control.

**Supporting information 2**

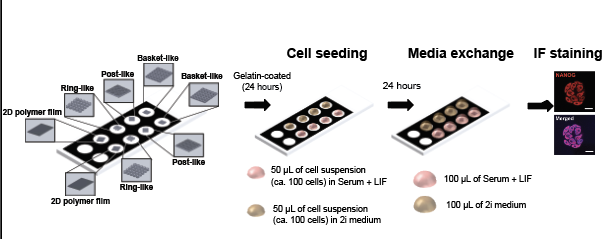

**Figure S2**. Optimized protocol for immunofluorescent (IF) staining. The microscope slide was washed with 70% ethanol and sterilized by UV-light (2.5 hours). Eight wells (two with polymer foil and six with ring-like, basket-like and post-like microscaffold arrays) were then coated by 0.2 % gelatin (24 hours). Cell seeding was performed in two steps: (1) placing a 50 μl medium droplet per well; (2) injecting 1 μl with ~100 cells above polymer foil or 3D microscaffold array. Afterwards, the microscope slide was placed in a 10 cm Petri dish with 3 ml of PBS to avoid medium evaporation. Media was exchanged after 24 hours. Finally, IF staining was performed (48 hours after seeding). Before imaging, a droplet of the mounting media (10 μL) was placed in the well, covered by a glass coverslip (170 m thick) and sealed.

**Supporting information 3**

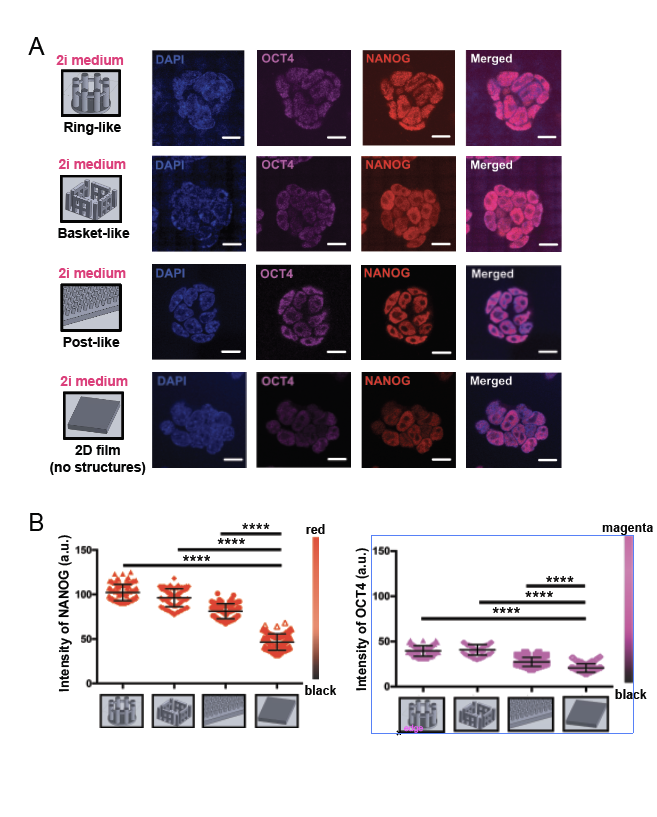

**Figure S3.** Representative immunofluorescent images of NANOG (red) and OCT4 (magenta) expression in mESCs grown in 3D microscaffold arrays and on 2D solid polymer film under 2i conditions. DAPI was used as a nuclear counterstain. Scale bar is 10 µm (A). Quantification of immunofluorescent intensity for the NANOG (red) and OCT4 (magenta) at the single nuclear level in mESCs cultured in three 3D microscaffold arrays (ring-like, basket-like and post-like) and on 2D solid polymer film under 2i conditions (B). Number of counted nuclei n=100. Immunofluorescence staining was performed in triplicate. P<0.0001 (Mann–Whitney test).

**Supporting information 4**

The generated GWL file with a programming code segment is presented below:

*% File generated by DeScribe 2.2.1*

*DefocusFactor 0.63 % refer to the manual for more details*

*% Writing parameters*

*GalvoScanMode*

*ContinuousMode*

*PiezoSettlingTime 100*

*GalvoSettlingTime 2*

*% System Initialization*

*TiltCorrectionOff*

*ResetInterface*

*% Field Parameters*

*XOffset 0*

*YOffset 0*

*ZOffset 0*

*% PiezoGotoX 0*

*% PiezoGotoY 0*

*PowerScaling 1.0*

*LaserPower 100*

*ScanSpeed 40000*

*Include slicer output include Ring-like microscaffold array_100_data.gwl*

The code was then loaded for direct laser writing process by the control software NanoWrite for manufacturing.
